# Deep-mutational scanning libraries using tiled-region exchange mutagenesis

**DOI:** 10.1101/2025.09.05.674540

**Authors:** Kortni Kindree, Claire A. Chochinov, Keerath Bhachu, Yunyi Cheng, Amelia Caron, Molly McDonald, Zaynab Mamai, Alex N. Nguyen Ba

## Abstract

The analysis of gene function frequently requires the generation of mutants. Deep-mutational scanning (DMS) has emerged as a powerful tool to decipher important functional residues within genes and proteins. However, methods for performing DMS tend to be complex or laborious. Here, we introduce Tiled-Region Exchange (T-REx) Mutagenesis, which is a multiplexed modification of the EMPIRIC mutagenesis approach. Self-encoded removal fragments are cloned in parallel in non-overlapping gene locations and pooled. In a one-pot reaction, oligonucleotides are then swapped with their corresponding self-encoded removal fragments in bulk using a single Golden Gate reaction. To aid in downstream phenotyping, the library is then fused with unique DNA barcodes using the Bxb1 recombinase. We demonstrate this approach and its optimizations, to show that it is both easy to perform and efficient. This method offers simple and expedient means to create comprehensive mutagenesis libraries.

**ARTICLE SUMMARY:** Researchers in the field of molecular genetics frequently investigate protein function via mutations. Here, we present a rapid deep-mutational scanning methodology called Tiled-Region Exchange Mutagenesis that can comprehensively create mutant libraries. The approach is a multiplexed version of the established EMPIRIC mutagenesis approach and retains its simplicity. We show optimizations of the in vitro reactions to reduce cost and improve robustness. This method will be useful to researchers seeking to perform deep-mutational scanning on any gene of interest.

## INTRODUCTION

In the field of genomics and proteomics, assessing the functional impacts of mutations has been a crucial area of research. Efficient strategies for site-directed mutagenesis have allowed improved understanding of catalytic enzymes(1), binding interfaces(2), and regulatory elements(3). Standard methods for site-directed mutagenesis frequently assess mutations at specific sites to specific residues and therefore only offer a small glimpse of possible mutations on a gene of interest(4). In proteins, a less biased method is alanine scanning(5), which aims to decipher the functional contribution of each native amino acid in a protein. More recently, alanine scanning has been supplanted by deep-mutational scanning (DMS) where the activity of each possible amino acid is compared at each position in a protein(6). These screens offer an extremely rich view of the functional and evolutionary constraints on proteins as amino acids of similar biochemical properties can be leveraged to obtain deeper insight on functional residues(6, 7). However, one major challenge with obtaining these improved mutagenesis screens is that deep-mutational scanning is far less accessible to current molecular biology labs due to cost and difficulty of production(8). This limitation has precluded the genome-scale analysis of variants.

To capture a complete set of comprehensive variants for a gene of interest, the DMS approach must be unbiased and provide a saturated library pool(9, 10). Ideally, every possible amino acid change across the target gene or region should be present, encompassing all variants identified through evolutionary comparisons or variant screening and large-scale sequencing efforts. Moreover, each variant created in the pool should be equally represented before having undergone any phenotypic assays. This approach must also have the capability of being performed in bulk, using high-throughput screening techniques, while being scalable to any desired gene size.

Several methods have been previously developed for generating variant libraries to assess mutations in bulk including parallel site-directed mutagenesis (SDM)(11), error-prone PCR (ep-PCR), Saturated Programmable Integration of Elements (SPINE)(12), and various oligonucleotide-based approaches (Kunkel(13), PFunkel(14), PALS(15), POPcode(16), and EMPIRIC(17)). These oligonucleotide-based approaches have become more popular in recent years due to the rapidly decreasing cost of oligo pool synthesis. While these methods all have their advantages, to our knowledge no existing method is both easy to perform and generates all possible single amino acid variants or missense mutations in a gene, while simultaneously ensuring that each plasmid copy contains only one single amino acid variant, and that the wildtype sequences are minimized in this library.

To address these current limitations, we have developed a simple, novel DMS approach that multiplexes the EMPIRIC methodology, called Tiled-Region Exchange (T-REx) Mutagenesis (Fig. 1). T-REx aims to achieve a high mutational efficiency while multiplexing the workflow within a streamlined one-pot reaction, for the purpose of creating a library of evenly represented single amino acid mutations along a gene of interest. This approach also aims to increase throughput, while decreasing the off-target mutation rate, to improve scalability for variant effect mapping and functional interpretation. Here, we outline our novel approach, its advantages, and implications as an integral DMS approach that generates a variant library enriched in mutations which may inform biomedical research and be relevant to disease.

**Figure 1:**
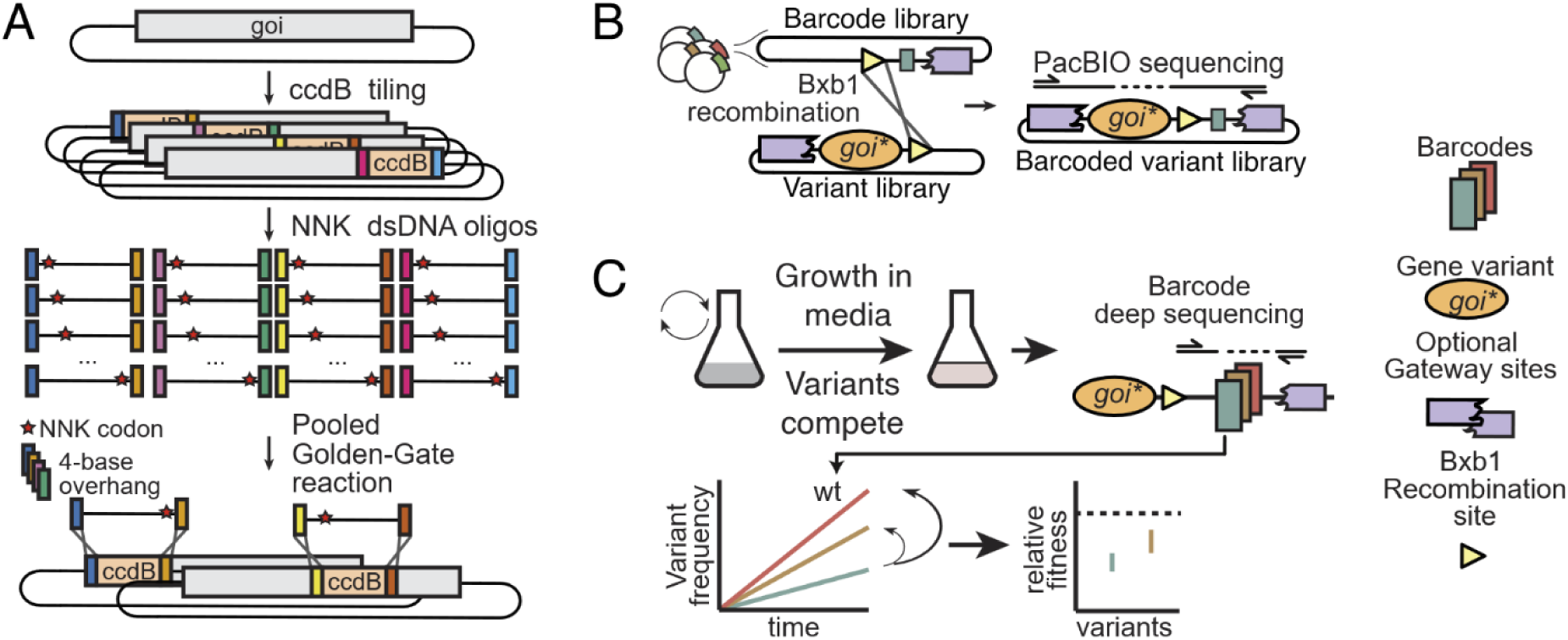
**(A)** The gene of interest (goi) is split into non-overlapping tiled regions with unique inward-facing BsaI sites. The native sequence is temporarily replaced with a *ccdB* negative selection cassette, and then swapped for the desired dsDNA mutagenic oligo containing NNK codons at each position in a Golden Gate reaction with 4-base overhangs. **(B)** The variant library and barcode plasmid library are fused in a Bxb1 recombinase reaction and can be sequenced to associate the mutation to their unique barcode. **(C)** To complete a selection assay on the variant library, variants can compete for growth and their relative fitness can be obtained by sequencing the abundance of each barcode before and after selection.

## MATERIALS AND METHODS

### In silico data-assisted design

To multiplex the EMPIRIC mutagenesis method, self-encoded removal fragments (SERFs) are cloned in non-overlapping locations (or tiles) in a gene in parallel, and then subsequently pooled followed by exchange with desired fragments using simultaneous Golden Gate assembly in a one pot reaction, where T4 ligase and BsaI restriction endonuclease work in tandem for scarless ligation reactions over non-palindromic overhangs. Reaction fidelity is ensured by the specific overhangs used in this Golden Gate reaction, but minimizing off-target ligations is still desirable by choosing tile locations that produce overhangs with minimal crosstalk.

To select these tile locations, we took inspiration from the Data-optimized Assembly Design process that was developed for simultaneous gene assembly of 52 fragments(18, 19). We begin by randomly choosing tile boundary locations and calculating an objective function, which scores the probability of on-target assembly, the probability of off-target assembly, the presence of palindromes, the presence of vector-only relegation, and the tile-size variance. Tile positions are then randomly moved in a local region and positions that maximize this objective function are maintained. After a few iterations, we obtain sets of tile locations with compatible overhangs that can be assembled in one-pot. A script implementing this algorithm can be obtained at: https://github.com/annb-lab/TRex.

### Library generation and cloning

#### *ccdB* entry vector cloning

Tiled *ccdB*-containing plasmids were generated with parallel PCR reactions, where the *Escherichia coli* plasmid containing a gene of interest was first amplified to exclude the tile region. Each of these PCR reactions were performed separately, so that there was an amplified fragment for each tile. The primers used for these PCRs contained a complementary overhang to the *ccdB* PCR fragment, which were then cloned with Gibson Assembly(20) in a 10 µl reaction by combining equal amounts of the *ccdB* fragment and backbone PCR fragment. Reactions were then incubated at 50 °C for 1 hour on the thermocycler. 1 µl of this assembled product was transformed into *ccdB* Survival 2 competent cells (Invitrogen), which allow for the otherwise toxic *ccdB* fragment to be propagated.

### Oligo extension

Oligos were ordered from IDT (Integrated DNA Technologies) as standard desalted oligos, as Ultramers, or as oPools (oligo pools from IDT, which use the Ultramer synthesis platform). They were then resuspended in 10 mM Tris, 0.1 mM EDTA pH 8.0 to 10 µM. Conversion of the single-stranded oligonucleotide to double-stranded DNA and preparation of the *E. coli* library was performed as in previously with small modifications (which will be described in their respective sections)(21). In some trials, gBlocks would substitute for these Ultramers, with specific introduced mutations, for a site-directed mutagenesis approach (three to four mutations in different tiles can be ordered as a single 500 bp gBlock) in a more cost effective and simple manner.

### NEBridge and transformation

Golden Gate assembly(22) was performed using a cycling method with a modified reaction buffer that shows higher cloning efficiency. We usually used about 500 ng of *ccdB*-entry vector, and 1 µl of purified dsDNA oligonucleotide in a 20 µl reaction. We used the following cycling protocol: 1) 37 °C for 1 minute, 2) 16 °C for 1 minute, 3) GOTO 1, 60 times, 4) 50 °C for 5 minutes, 5) 12 °C for infinite time. We used a 5x assembly buffer: 200 µl of 10x Cutsmart buffer, 20 µl of 100 mM ATP, 20 µl of 1 M DTT, 60 µl of propylene diol, 100 mg of PEG 8000 and water to 400 ul. Our 20x enzyme mix was: 10 µl T4 ligase (2000 U/µl), 30 µl BsaI-HFv2 (20 U/µl). The product of the cycling reaction can be directly transformed into homemade chemical competent cells(23), and we routinely obtain 10^5^-10^6^ cfu. from this transformation.

### Bxb1-integrase fusion

To barcode the deep-mutational scanning library, barcodes were cloned in a plasmid containing kanamycin resistance and an R6K origin of replication, which requires a strain containing the Pir element for propagation(24). Barcodes were cloned as in previously, except using strains BAN004 (DH10B, *uidA*::pir116) and BAN005 (ccdB Survival 2, *uidA*::pir116) for plasmid propagation. The plasmid also contained a Bxb1 attP site, while the variant library, containing ampicillin resistance and a pUC origin of replication, contained a Bxb1 attB site(25). A Bxb1 fusion reaction was prepared as follows: 400 ng of the plasmid containing the mutagenized library, 200 ng of the barcode library, 2 µl of 5x Bxb1 buffer (250 mM Tris pH 8, 250 mM KCl, 375 mM NaCl, 5 mM EDTA, 500 ug/mL BSA, 25% PEG 8000), 1 µl of Bxb1 integrase enzyme (at approximately 0.5 mg/mL) and topped with water to 10 ul. The reaction was then incubated at 37 °C for 2 hours in a thermocycler. Gel electrophoresis verification of the fusion reaction indicated that the reaction is essentially complete within 2 hours at 37 °C. The reaction was then transformed directly into chemical competent *E. coli* cells and recovered in LB+Amp+Kan to select for cells that received the fused plasmid only.

### Purification of Bxb1 integrase

Purification of the Bxb1 integrase was performed using a Bxb1 integrase construct that was C-terminally tagged with a 6xHis. This construct was expressed in BL21(DE3) *E. coli* cells that were grown in ZYM505 medium(26). The cultures were induced with IPTG at 0.25 mM at mid-log phase (OD₆₀₀ ∼ =0.4–0.8) and incubated overnight at 18°C with shaking. Cells were then harvested by centrifugation and resuspended in lysis buffer (20 mM Tris (pH 8), 1 M NaCl and 5% (v/v) glycerol). After cell lysis by sonication and centrifugation to remove cell debris, the clarified lysate was loaded into a nickel-nitrilotriacetic acid (Ni-NTA) resin column, which was pre-equilibrated with lysis buffer + 10mM Imidazole. The column was then washed with the same lysis buffer + 10mM Imidazole, to remove excess non-specific binders and outcompeting weakly bound proteins, before eluting the 6xHis-tagged Bxb1 integrase with lysis buffer + 250 mM Imidazole. The eluted protein then underwent a buffer exchange to completely remove the imidazole and concentrate the protein to 1 mg/ml. The purified protein was then diluted with equal volumes of glycerol (final 50% v/v glycerol), reaching a final concentration of ∼8.5 µM and stored at –20 °C. We did not remove the 6xHis-tag from the Bxb1 integrase protein since the tag is small (∼0.8 kDa) and has not been seen to interfere with the robust, high-yielding Bxb1 reaction efficiency.

### Yeast transformations

After barcoding the library, the purified plasmid pools can be transfected into a model organism of choice. We used a standard Lithium Acetate (LiOAc) and 50% PEG protocol to transform our libraries into yeast(27). An overnight culture of the desired yeast strain was grown to saturation in YPD (2% peptone, 1% yeast extract, 2% (w/v) glucose), and 150-300 µl was transferred into 5 ml of fresh YPD and grown for 4.5-5 hours at 30 °C until an optimal OD600 of 0.4-0.6 was reached. The cells were then pelleted, washed, and the following were added on top of the cells: 240 µl of 50% PEG 3350, 36 µl of 1 M Lithium Acetate, 50 µl denatured Salmon Sperm DNA (2 mg/mL, boiled 5 min, snap cooled on ice), 10-20 µl of PmeI digested mutagenized barcoded library. The PmeI digestion was done to release the mutagenized fragment with regions of homology for integration in the yeast genome. It is specific to our experiment and other means of transfection are possible. The reaction was then vortexed until homogenized and left to incubate in a 42 °C heat bath for 1 hour. The cells were then pelleted, the supernatant was removed, and the pellet was resuspended in 1 ml of ddH2O before being plated on standard dropout plates and incubated for 2-3 days at 30 °C.

### Sequencing library preparation

#### gDNA extraction

After growing the transformed yeast pools to saturation, the genomic DNA was extracted using the following gDNA extraction protocol. 1-3 mL was spun down and the supernatant was removed. The cell pellet was resuspended in 100 µl of yeast lysis extraction buffer (5 mg/mL Zymolyase 20T (100 U/mL), 100 mM Sodium Phosphate buffer pH 7.4 (0.5 M Na2HPO4 and 0.5 M NaH2PO4), 10 mM EDTA, 0.5% SB3-14, 200 µg/mL Rnase A, 1 M Sorbitol, 20 mM DTT, stored at -20 °C) and placed at 37°C for 30 minutes or more until complete lysis. After lysis, 400 µl of lysis/binding buffer (100 mM MES pH 5, 4.125 M Guanidine thiocyanate, 25% isopropanol, 10 mM EDTA) was added to the tube, and vortexed until all precipitates were dissolved. The tube was spun down for 30 seconds if unlysed cells remained. The supernatant of the tube was passed onto a standard miniprep silica column and spun for 30 seconds to pass the supernatant through the column. 1x wash with 400 µl Wash buffer 1 (10% GuSCN, 25% Isopropanol, 10 mM EDTA) followed by 1x wash with 600 µl 10 mM Tris/80% Ethanol was performed and spun for 30 seconds after each wash. The column can then be dried by spinning for 3 minutes at maximum speed. The gDNA was then eluted with 50 µl of elution buffer (10 mM Tris-HCl, pH 8.5), where the expected DNA concentration was around 20-30 ng/µl for a 1 mL culture. Success of the extraction was verified by agarose gel electrophoresis.

### PCR and sequencing

Unique barcodes linked with mutated codons can be sequenced using long-read sequencing or with Illumina in the case of short libraries. This can be done with PCR (necessary for libraries integrated inside model organisms), or by restriction digest followed by adapter ligations according to the recommendations of the sequencing platforms. Our FKBP1a library was sequenced on a MiSeq v2 500-cycle kit using PCR protocols as previously described in a previous study(21).

### Barcode association bioinformatics

#### Read extraction of genes and barcode sequences

Reads were de-multiplexed from inline indexes and barcodes were analyzed as in a previous study(21). To extract the gene from the reads, 20 bp anchors corresponding to the promoter and terminators were used, allowing 2 mismatches per anchor. The barcode was similarly extracted, removing fixed bases included in the barcode to prevent BsaI restriction enzyme sites.

### Barcode clustering

Sequenced barcodes contain a mixture of barcodes with and without sequencing errors. Assuming that most sequencing reads have no base calling mistakes on the barcodes, we can perform clustering and error correction to obtain a final set of barcodes. This was performed essentially as in (21), by sorting barcodes by their counts, and error correcting barcodes starting from least common to most common, using a threshold of 2 Hamming distance. Finally, barcodes that were observed fewer than 10 times in the whole sequencing library were removed from further analyses.

### Mutation association

To associate mutations within the gene to a barcode, we tabulated for each barcoded read the list of mutations detected. Each read can contain, in principle, four different types of mutations: 1) the correct codon that was mutated by Tiled-Region Exchange mutagenesis, 2) an oligo synthesis mistake that is associated with the mutated codon, 3) a sequencing error due to basecalling, and 4) a sequencing error due to PCR chimeras. Because oligo synthesis errors will occur at the same frequency as the desired mutated codon (forming a mutation set), we take advantage of the Apriori algorithm (28) to identify the most common set of mutations for each barcode at the nucleotide level. Given a total list of possible mutations associated with a barcode, the Apriori algorithm can return, for each possible set of mutations, the most common to least common sets. Sequencing errors occur usually as singletons and can be filtered out by a simple frequency threshold. However, PCR chimeras occur during the amplification reaction and can therefore appear in several reads. To investigate the effect of PCR chimeras in our analysis, we first cumulated the frequency of the most common mutation set and the second most common mutation set for each barcode (assuming that the most common mutation set is the real mutation associated with a specific barcode). We found that the most common mutation set was usually found at over 30% of the reads, while the second most common mutation set was usually below 10% of the reads. Thus, implementing a mutation set threshold of 30% and a minimum read count of 10, both accelerates the Apriori algorithm and likely returns true barcode-mutation associations.

If two mutations were found at over 30% of reads, this would indicate barcodes that are shared between different mutations. These were discarded from our analysis, as well as barcodes that were associated with an indel.

## RESULTS AND DISCUSSIONS

### Efficient and comprehensive deep-mutational scanning libraries using tiled Golden-Gate assembly

There is no shortage of approaches for systematically mutagenizing genes of interest on a plasmid. The conceptually simplest approach to do this is by performing standard site-directed mutagenesis (SDM) at every desired position, however this is extremely laborious(11). Higher throughput methods have used mutagenic oligonucleotides and a polymerase, or error-inducing enzymes (such as error-prone PCR(29) or fusion of T7 RNA polymerase with AID(30)). All these approaches, however, require fine-tuning the Poisson rate of introduced mutations and cannot guarantee that all constructs contain a single mutation (and not zero or more than one). In contrast, the EMPIRIC (Extremely Methodical and Parallel Investigation of Randomized Individual Codons) method is an oligonucleotide-based approach that clones the designed oligonucleotides directly with the use of a ligase (17) and thus falls into a different class of mutagenesis techniques that can guarantee the final product. In EMPIRIC, plasmids receiving the oligonucleotide cassettes have a Self-Encoded Removal Fragment (SERF) flanked by inverted BsaI restriction sites(17). As such, plasmids can be treated enzymatically with BsaI to ligate oligo fragments flanked by sticky ends that have been designed to ligate at the desired position in the target vector. Where the EMPIRIC method falls short is that it only mutates a single region of a gene. However, it is trivial to imagine how the EMPIRIC approach can be parallelized to mutagenize a complete gene.

In light of this, the theoretical ideal approach would generate one and only one mutation, with high mutational efficiency in a one-pot reaction. This would allow the production of comprehensive libraries of all possible missense variants in a gene of interest. To this end, we developed Tiled-Region Exchange (T-REx) mutagenesis, which was greatly inspired by the EMPIRIC approach. Briefly, the approach is a *multiplexed* version of EMPIRIC where all the SERFs (referred to as ‘tiles’) are replaced with their corresponding oligonucleotides in a single reaction. The following four objectives were prioritized prior to the development of this method: ease of use, generation of all possible single-mutant variants, production of one and only one variant per clone, and minimization of unmutated sequences in the library.

To multiplex EMPIRIC and decrease unmutated sequences in the library, we made two major modifications to the protocol: 1) we include the toxic *ccdB* gene in the SERF, and 2) we use tiles that are positioned such that they contain unique and optimized overhangs that can be used to clone synthetic mutagenizing oligos in a single reaction. The addition of *ccdB* in the SERF virtually guarantees that only cells that have undergone a successful tile-exchange will be viable(31), thus decreasing the frequency of clones that maintain the WT, unmutated sequence. The unique optimized overhangs enable the entire gene to be mutated in a single reaction vessel as these minimize off-target ligations of oligonucleotides. Finally, to aid in downstream phenotyping of mutants, we introduce an approach to add unique barcodes to the mutagenesis libraries using the Bxb1 recombinase.

A brief overview of the workflow of T-REx is as follows: parallel cloning of SERFs in a gene of interest on a plasmid, converting designed oligonucleotide pools to dsDNA, one-pot cloning of all the oligonucleotides with a plasmid pool of SERF-containing genes (Fig. 1A), followed by one-pot fusion with a unique barcode (Fig. 1B). The final library can be introduced in a model system of choice for further phenotyping (Fig. 1C).

### Automated and optimized design of tile locations

One critical component of T-REx is the initial insertion of SERFs. Simultaneous cloning of oligonucleotides that potentially encode different tile sequences requires that their ligation and exchange with corresponding SERF is highly specific. Here, a trade-off between the number of tiles and the length of the cloning oligonucleotide must be considered, as longer oligonucleotides are more costly and may have a higher rate of synthesis errors (Ultramers from IDT are expected to yield full length oligos at 50% of the unpurified products at 140 bp (32), while having more tiles increases the complexity of the reaction (notwithstanding the increased number of SERFs that must be cloned in parallel).

To explore these constraints, we first verified whether the length of mutagenic oligonucleotides is a limiting factor in T-REx mutagenesis. We cloned SERFs inside regions of different lengths into the eforRed pink-producing chromoprotein(33), and performed a single oligonucleotide exchange, counting resulting colonies for cloning efficiency and yield. A successful reaction yields pink colonies, while oligonucleotide synthesis errors (which are frequently indels) would yield a white colony. Across a variety of overhangs and a large range of oligonucleotide sizes (from <60 bp to 175 bp), using standard desalted oligos or using Ultramers from IDT (for longer oligos), we found no prohibiting differences in correct assembly or colony counts after *E. coli* transformation (Fig. S1). Others have also found similar rates of full-length synthesis with Ultramers from IDT (34) and as will be described later, even tiles of 120 bp (40 amino acids) that require oligonucleotides of about 180 bp can be mutated successfully with minimal errors (∼5% indel rate, and ∼5% wrong base synthesized). Therefore, contrary to our original assumption, the length of the oligonucleotides or the synthesis platform is not a major limiting factor for EMPIRIC and the requirement for semi-accurate synthesis on both 5’ and 3’ ends of the purchased oligo is sufficient to ensure high-efficiency cloning. While shorter oligos may be more cost-effective and accessible, they do not strongly influence the reaction efficiency, and thus the trade-off for a comprehensive mutagenesis library is simply the cost of oligonucleotides and the total number of SERFs. In a practical sense, the span of a tile can therefore be about 40 amino acids.

To reduce cost, we further explored the number of bases needed for cleavage close to the end of DNA fragments for the BsaI restriction endonuclease. According to NEB, BsaI can cleave fragments when the recognition sequence is 1 bp from the end of a double-stranded fragment, but they recommend 6 base pair for Golden Gate assembly(35). We thus varied the number of bases from 0-20 bp past one of the two BsaI site in the oligo (one site is necessarily longer due to the need for a primer during conversion of ssDNA oligos to dsDNA, see Methods) and tested the efficiency of the reaction using the same colorimetric assay as previously discussed except that we mutagenized a plasmid containing the amilCP blue-producing chromoprotein. In our hands, all extensions, including 0, 1, 2, 3, 4, 5, 6, 7, 8, and 17 bp, supported a robust assembly with high colony counts (Fig. 2A and Fig. S2 for similar experiment using the amilOrange chromoprotein).

**Figure 2:**
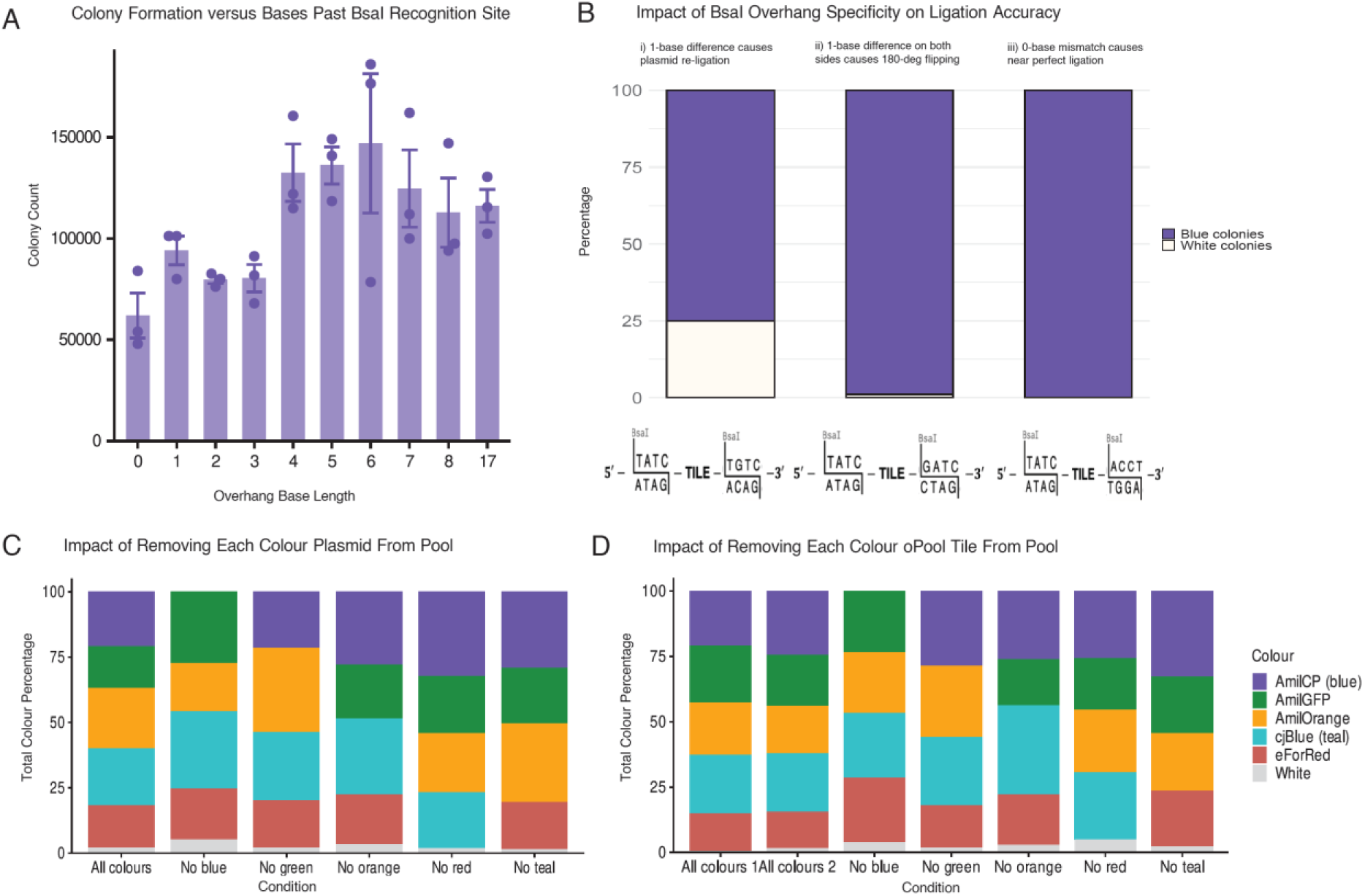
**(A)** Barplot depicting colony formation vs the number of overhang bases past the BsaI recognition sites, with three replicates per base length. **(B)** Stacked barplots indicating the overhang specificity and its impact on ligation accuracy. **(C-D)** Stacked barplots indicating colony counts grouped by chromoprotein identity (five chromoproteins each represented by a different colour). **(C)** depicts the difference between all colours included in the pool, versus omitting each colour plasmid individually from the pool, while **(D)** shows the colony counts of the pool when omitting each oPool oligo individually from the pool of oligos.

Upon testing many different overhangs, we encountered one case where an increase in white colonies was observed and traced this to a case where overhangs differing by one base could promote re-circularization of the plasmid without the oligonucleotide. While this is a common occurrence in some restriction-endonuclease cloning workflows, it was not expected to occur based on the measured fidelity of T4 DNA ligase (5-GGAA ligated to 3-CCCT was seen approximately 0.1% of the time in NEB’s screen(18)). Despite this, we observed about 25% white colonies in this reaction (Fig. 2B). Thus, care must be taken when designing tile locations to minimize these off-target ligations.

Our previous results suggest that T-REx could be used as a comprehensive mutagenesis technique that can mutagenize whole genes with minimal constraints. Thus, to aid in developing this methodology, we developed a script that can automate the cloning process and choose optimal tile locations, inspiring ourselves from data-optimized assembly design. The efficiency and fidelity of T4 DNA ligation on all possible 4 bp overhangs was previously measured by NEB(18), and serves as a guide for optimized tile locations. The number of tiles is chosen by the user (taking into account the total cost of assembly), and all oligonucleotides, including mutagenic oligo pools, are returned in a simple output.

While it has been routine in synthetic biology labs to remove BsaI sites from genes for cloning purposes(36), it is also necessary for deep-mutagenesis to consider that sets of random nucleotides can generate a *de novo* BsaI restriction sequence within the mutagenized region. In our lab, we usually perform degenerate NNK mutagenesis (though NNN or NNS is also possible) to randomly generate all 20 amino acids at a single site, while reducing redundancy and the number of stop codons. In certain instances (for example NNK CTC), a spurious BsaI site can be produced. In these cases, the script automatically attempts to further mutagenize surrounding bases to preserve the coding amino acid (for example NNK CTC to NNK CTT) or will resort to NNS mutagenesis. One advantage of data-optimized assembly design is that it can easily incorporate further constraints in the design, and so other restriction endonuclease sites can also be automatically screened and removed in a similar manner.

### Tiled-Region Exchange mutagenesis can be performed as a one-pot reaction

One strength of T-REx is that, by choosing unique overhangs where off-target ligations have been minimized, the reaction might be performed in one-pot. Though not necessarily required for successful and comprehensive mutagenesis, this may improve throughput and cost as long as the reaction remains specific. For example, the list price for IDT oligo pools in small scale is 5 cents per base, while it is 1 cent per base at very high scale(32). To showcase the specificity of this one-pot reaction, we cloned SERFs into 5 different chromoproteins. Single-stranded oligos that ‘repair’ these SERFs were ordered and pooled prior to conversion to dsDNA, and a one-pot Tiled-Region Exchange mutagenesis reaction was performed. The tile locations were chosen such that incorrect assembly (an oligo swapped with the wrong SERF) would yield an unpigmented colony. Under ideal conditions, we would expect approximately equal cloning efficiencies for all five chromoproteins and thus observe an equal proportion of coloured colonies. The results of this experiment are shown in Fig. 2C-D, suggesting that the reaction is highly efficient and specific, depicting <1% of colonies having been incorrectly ligated and appearing white, with each colour appearing at a similar frequency.

To further highlight the specificity of this reaction, we performed the same experiment but this time omitting either one plasmid entirely (Fig. 2C) (and thus having too many ‘repair oligos’) or omitting one swapping oligonucleotide (Fig. 2D) (and thus having one plasmid with a SERF that cannot be successfully cloned). In neither of these cases did we observe a detrimental effect. As shown in Fig. S3, all 4 of the other chromoproteins were properly ligated with their exchange oligos and there was no excess of white colonies. As expected, no colonies produced the omitted intact chromoprotein.

### Barcodes can be attached to the mutagenic library using recombinase-mediated fusion

In the EMPIRIC approach, the mutagenized fragment can be sequenced directly for phenotyping purposes using standard short-read sequencing. However, when mutagenizing whole-genes in one-pot reactions, this approach cannot be guaranteed to sequence the intended tile and long-read sequencing typically do not offer the throughput required for phenotyping thousands of variants effectively. Previously, it has been shown that short, unique DNA barcodes can be linked to variations of interest and used to phenotype libraries effectively with short-read sequencing(37).

In many deep-mutational scanning protocols, DNA barcodes are incorporated into the mutagenized library using PCR, by incorporating random nucleotides in the primers and cloning of the amplified product (38), or by direct ligation (39). Due to the large number of random bases in these designs, this approach virtually guarantees that barcodes do not associate with more than one gene fragment, enabling high fidelity in the following analyses. However, if the PCR product that is amplified with the barcodes is the mutagenized library, then the final libraries may contain PCR chimeras and contain more than one variant. To remedy this, mutagenized libraries can be isolated by restriction enzymes and ligated on a barcoded PCR product of the desired plasmid backbone.

Here, we chose a different strategy where barcode libraries can be sequenced and characterized beforehand, effectively generating subsamples of the random nucleotide space. These barcodes can be sequenced at high depth, aiding in future long-read mutation association sequencing. This library, however, must be fused, or linked, to the mutagenized genes. To do this, we designed our mutagenesis to be performed on a plasmid containing a Bxb1 attB(25) (bacterial attachment) integration site and ampicillin resistance. On another plasmid, a barcode library is cloned on an attP-containing (phage attachment) plasmid(25) with an R6K origin of replication with kanamycin resistance(24). Bxb1 serine-integrase can thus fuse both plasmids *in vitro* on the conserved “GA” dinucleotide sequence within the attachment sites, yielding a barcoded deep-mutation scan library after transformation into standard DH5a cells and selecting on both kanamycin and ampicillin. The linkage between the mutagenized variant and the unique barcode can be determined by using any of the long-read sequencing technologies available and the use of Bxb1 integrase enables flexibility for downstream users that may wish to use the Gateway(40) system to clone their library onto different backbones while preserving this linkage. Thus, this approach allows the unique tagging, quantification, and identification of each variant in the sequencing analysis from complex multiplexed mixtures.

In our workflow, we have optimized this Bxb1 fusion reaction through several parameters and assessed these reactions on an agarose gel and by transformation. We first tested a range of temperatures, 20 °C, 25 °C, 30 °C, and 37 °C, where we noticed that the most complete fusion reaction occurred at 37 °C (Fig. 3A). We also tested various incubation times, at 37 °C, by periodically plating the reaction every 15-30 minutes and found that the number of colonies obtained was generally sufficient after two hours, which roughly corresponded to results from the agarose gel electrophoresis (Fig. S4). We also tested various salt concentration and additives (Spermidine, Bovine Serum Albumin (BSA), propylene glycol, polyethylene glycol (PEG), etc.) to the fusion buffer, where good fusion results occurred in the presence of at least 75 mM NaCl (Fig. 3B), 50 mM KCl and incorporation of 5% PEG 8000 (Fig. 3C and Fig. S5). Finally, to confirm fusion of both plasmids, restriction enzyme digestion was used to show a successful reaction (Fig. 3D).

**Figure 3:**
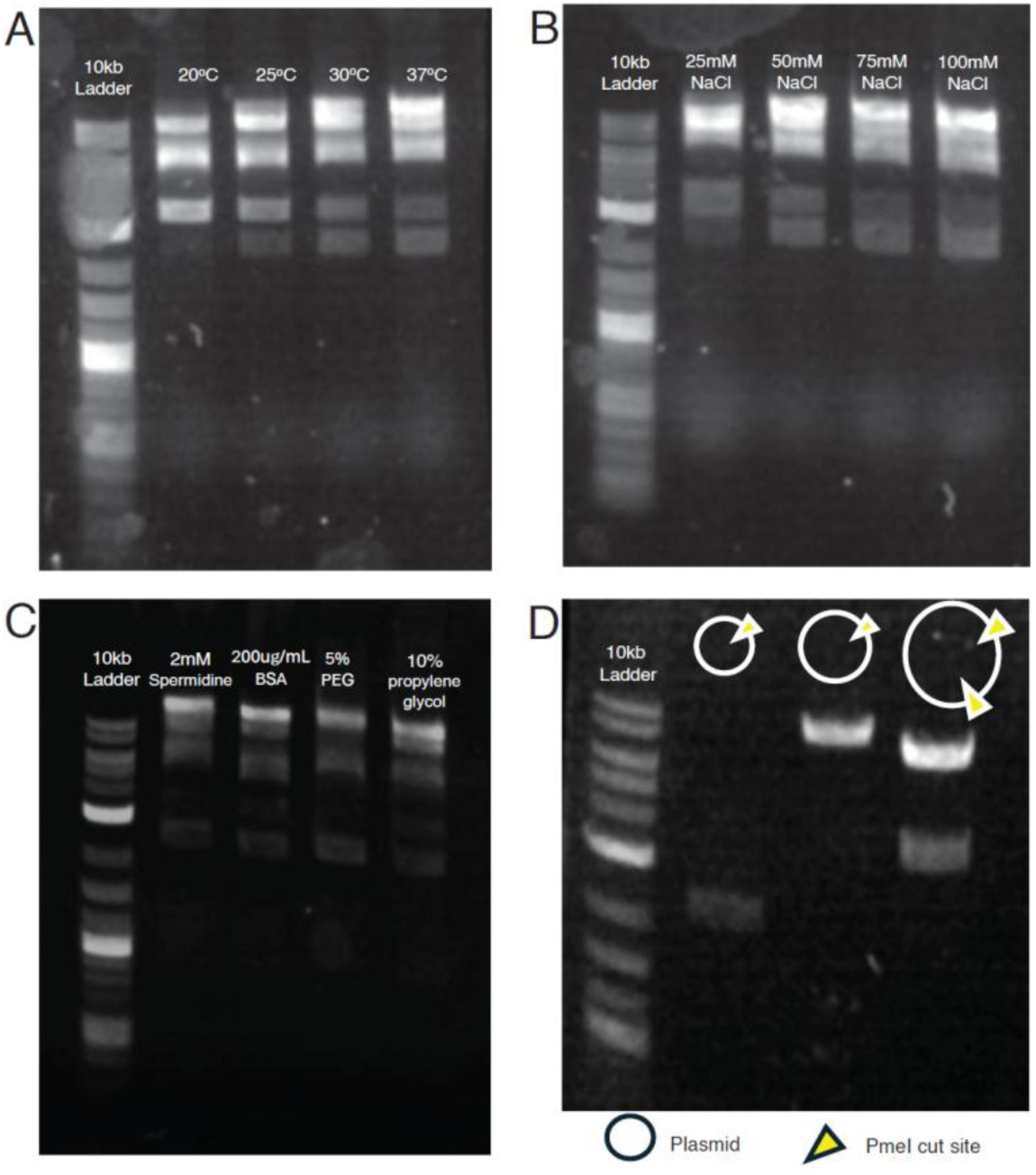
Optimization of Bxb1 integrase-mediated fusion between a linear R6K (oriγ) recipient vector and a donor fragment. A complete fusion reaction yields four diagnostic bands corresponding to the linear R6K backbone, the donor fragment, and the two recombination products and is consistent across each reaction. **(A)** Agarose gel of temperature series. Lane 1: DNA size ladder. Lanes 2-5: fusion reactions performed at 20 °C, 25 °C, 30 °C, and 37 °C, respectively for 1 hour. **(B)** Additive series. Lane 1: DNA ladder. Lanes 2-5: reactions supplemented with 2 mM spermidine, 200 µg/mL BSA, 5% PEG, and 10% propylene glycol, respectively. Reaction at 37 °C for 1 hour. **(C)** Salt tolerance series. Lane 1: DNA ladder. Lanes 2-5: reactions supplemented with 25 mM, 50 mM, 75 mM, and 100 mM NaCl, respectively. Reaction at 37 °C for 1 hour. **(D)** Validation of plasmid fusions using PmeI restriction endonuclease which confirms correct linearization of fragments. **Lane 1:** 10kb DNA ladder; **Lane 2:** PmeI digested barcode plasmid before Bxb1 fusion reaction; **Lane 3:** PmeI digested mutagenized plasmid before Bxb1 fusion reaction; **Lane 4:** PmeI digested fusion construct of barcode plasmid and mutagenized plasmid after Bxb1 fusion reaction. Band sizes are consistent with PmeI restriction site locations.

One drawback of our approach is that the barcode diversity is necessarily much lower than barcodes added by PCR or by ligation. Indeed, this diversity is limited exactly by the number of barcodes found in the barcoded R6K plasmid pools. However, several lines of evidence suggest that this disadvantage can be circumvented by simply having a modestly large number of barcodes. First, a barcoded R6K plasmid pool from our barcoding construction protocol usually contains about 250,000 barcodes as established from colony counts, far exceeding the number of variants found in most deep-mutational scan studies. For example, a gene of 1000 amino acids will have 32 000 possible single-NNK mutants, allowing about ∼8 barcodes per variant as biological replicates. We believe having fewer biological replicates (such as ∼8) is preferential to enable high-quality phenotyping as 1000 reads per barcode may be required to estimate barcode frequencies. Second, even if barcodes get associated with two different mutations, this should occur relatively infrequently as we will discuss in the next section. To assess this more generally, we reanalyzed the dataset from (21), which used the same barcode library to insert random barcodes into several yeast strains that contained a fixed known DNA barcode. Analyzing the total set of barcodes from the libraries yielded 188478 barcodes (from the expected ∼250000 from colony counts). This library was inserted into two starting yeast strains, with the first having 41207 barcodes and the second yeast strain having 28013 barcodes. Between the two yeast populations, the number of shared barcodes was 5227, which was close to the expectation of 6119. Thus, the number of barcodes that will be associated with the same mutations is directly controlled by the barcode library size, which can be built to be sufficiently large for most deep-mutational scanning studies. Finally, if this overlap is found to be too high, it is trivial to obtain more barcodes by constructing more barcoded R6K plasmid pools.

### Deep-mutational scanning libraries produced by tiled-region exchange mutagenesis are comprehensive

To confirm that Tiled-Region Exchange mutagenesis can be used for the production of comprehensive single-variant libraries, we produced a mutant library for the human FKBP1a gene, whose protein product binds to TOR in the presence of rapamycin to inhibit cell growth(41). The library was produced in two technical replicates and was integrated into the yeast genome at the benign *ho* locus, which contains loss-of-function mutations in the laboratory yeast strain(42). We then sequenced the barcoded libraries after yeast integration and sought to obtain several quality metrics that could confirm successful mutagenesis. The biological insights gained from performing comprehensive variant scanning on FKBP1a after selection will be described elsewhere.

We first verified whether NNK mutagenesis ordered as oPools had a nucleotide bias, which would result in a skewed amino acid representation. In standard desalted oligos, IDT offers ‘hand-mixing’ to ensure equal base representation, but this cannot be done at the scale of oPools. We found a modest synthesis bias with 30%G: 29%T: 24%A: 17%C. With NNK mutagenesis, this translated to an overrepresentation of glycine amino acids, and a decrease in histidine and glutamines. While proline was also reduced compared to expectations, there are more codons that code for proline than histidine, and as such the absence of prolines in the final mutagenesis library was not very pronounced. To rectify this bias, we suggest that oligos can be synthesized as both Cricks and Watson strand (so as to have MNN as well), however this will double the cost of making such mutagenesis libraries and it may be more cost-effective to transform the libraries at higher multiplicity if possible.

Finally, to show that the library can produce variants at every position, we cumulated sequencing reads with barcodes and their consensus mutations. In total, 14,476 barcode-mutation association pairs were obtained (7104 from replicate 1, and 7372 from replicate 2). As mentioned previously, only a small number of these barcodes were associated with oligo synthesis errors despite pooled oligo lengths of 140 bp to 179 bp: 588 contained a frameshift (4%) and 425 (3%) barcodes were associated with two mutations, either by oligo synthesis error or through fusing of the same barcode to two variants. To verify whether our barcode library was diverse enough, we found 269 barcodes present in both libraries that were associated with different mutations (close to the expectation of 259). Thus, the diversity of our R6K barcode library of ∼250,000 barcodes was relatively high enough to ensure only a small number of unusable barcodes.

We found on average 4.35 barcodes per NNK codon, with the median being 3. However, we observed a slight mutational bias in the first tile (Fig. 4A), presumably due to uneven plasmid mixing or exchange efficiency, our mutational coverage was 94.66% and a heatmap indicating positions with the number of barcodes representing the mutation is shown in Fig. 4B. Obtaining about 100 colony-forming units per mutational position appears sufficient to ensure high coverage of all variants. These results suggest that T-REx mutagenesis can be used for the production of comprehensive deep-mutational scanning libraries.

**Figure 4:**
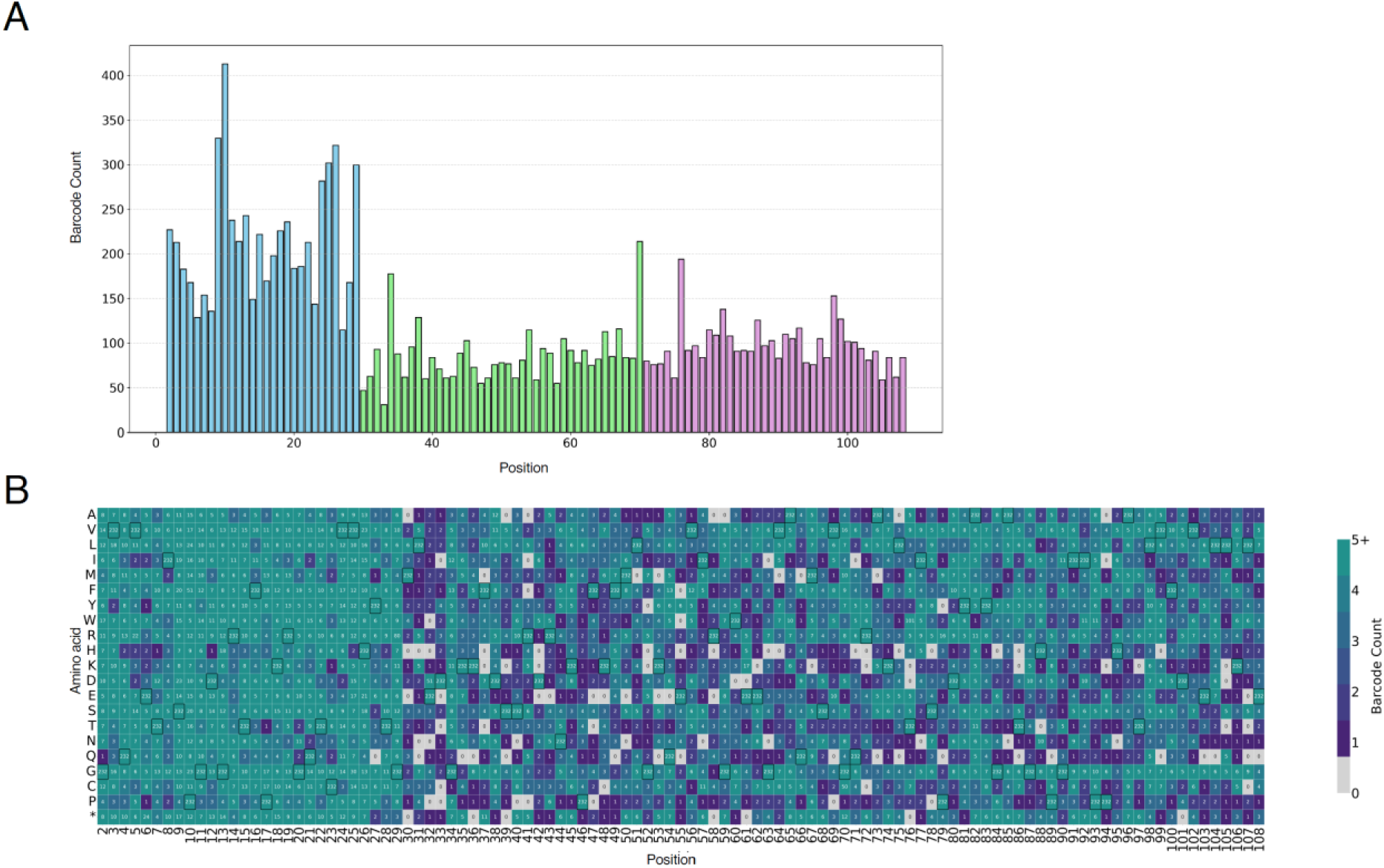
**(A)** Bar plot showing the number of barcodes per position along FKBP1A. The gene was divided into three tiles: Tile 1 (positions 2-29), Tile 2 (positions 30-70), and Tile 3 (positions 71-108) and a total of 13,463 barcodes-mutation pair were detected, distributed as follows: 6,279 in Tile 1, 3,743 in Tile 2, and 3,950 in Tile 3. Barcodes corresponding to variants without a single identifiable mutation or an indel were excluded from this analysis (1013 barcodes removed). **(B)** Heatmap of barcode frequency per mutation. Wild-type positions are outlined in black boxes.

## CONCLUSION

Tiled-Region Exchange (T-REx) mutagenesis combines the simplicity of the EMPIRIC approach and the throughput of other one-pot mutagenesis techniques. One caveat, however, is that the initial *ccdB* cloning stage must be done in parallel and can be laborious for large genes or for more systematic studies. Further, while T-REx can essentially guarantee one mutation per plasmid within the pool, it is often desirable to introduce multiple mutations per fragment. With T-REx, multiple variants can be introduced by designing oligonucleotides containing several mutations, however the user is constrained to a small region within a tile. In principle, this can be rectified by assembling several oligonucleotides simultaneously (in a three- or four-piece Golden Gate assembly), however this can only produce combinatorial libraries. In cases where single variants must be observed under different gene backgrounds, it may be preferable to combine T-REx with error-prone PCR or with other mutagenic techniques.

During the writing of this manuscript, another study showcased a similar assembly technique for deep-mutational scanning(43). Our approach differs in a few ways, namely by using *ccdB* during one of the reaction steps, and by allowing pooling of the complete reaction during assembly. In contrast, one particular strength of that assembly method is that barcodes are programmed during mutagenesis, which avoids the need for long-read sequencing and can even bypass sequencing to associate barcodes with mutations. Nevertheless, as deep-mutational scanning becomes more accessible, there will be a rise of new methodologies that increase efficiency, cost, and ease of use. New developments in this area will enable combining the strength of different established methodologies to ultimately give flexibility to researchers in this field.

Despite the limitations of Tiled-Region Exchange mutagenesis, our lab has leveraged the relatively straightforward methodology to train undergraduate students with minimal molecular biology experience, and we anticipate that its ease of use will be useful for labs seeking to produce routine comprehensive variant libraries without much trial and error. Finally, Our FKBP1A library cost about $600 CAD for the oligos required to produce it, while a pre-made variant libraries from Twist Bioscience was quoted at $5400 USD. Thus, the cost of making libraries in-house is also relatively competitive compared to commercial products.

## DATA AVAILABILITY

Strains and plasmids are available upon request. Raw sequencing data has been submitted to Sequence Read Archive. BioProject ID: PRJNA1316630. Code is available on https://github.com/annb-lab/TRex.

## ACKNOWLEDGMENTS

KK, CC, KB, and ANNB acknowledge funding from the Canadian Institute of Health Research (CIHR: Canada Graduate Scholarship-Master’s, OGB-185738). ANNB also acknowledges funding from NSERC (RGPIN-2021-02716), Connaught, and the OVPRI at UTM. We also acknowledge Fritz Roth for input during the development of this method. Finally, we acknowledge The Centre for Applied Genomics sequencing center for technical help during the sequencing of the libraries.

## Notes

https://github.com/annb-lab/TRex

### Competing Interest Statement

The authors have declared no competing interest.

### Summary of Updates

Figure 2 revised; Supplemental files updated; Discussion updated

