## Supplemental Figures S1-S5 for "Deep-mutational scanning libraries using tiled-region exchange mutagenesis"

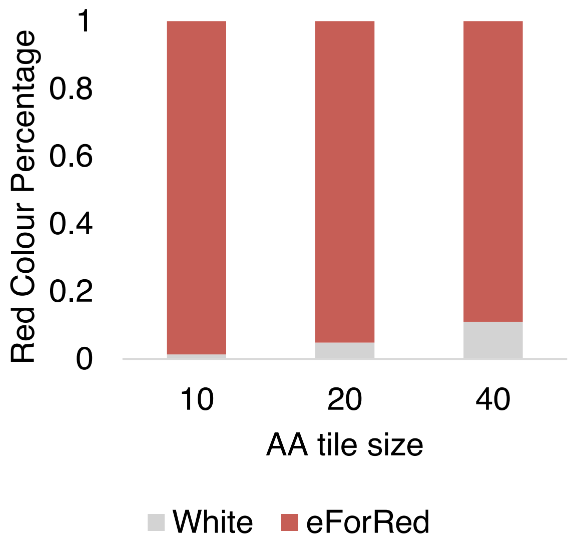


**Figure S1:** Effect of increasing oligo length on the proportion of successful reactions as assessed by red chromoprotein formation after transformation.

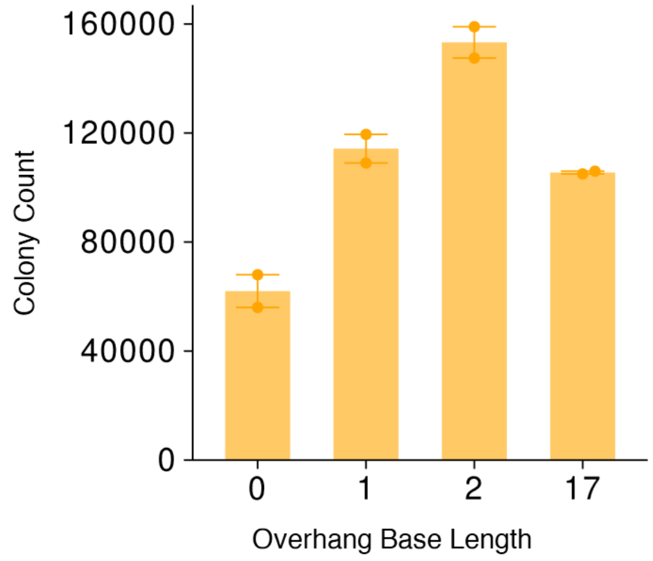


**Figure S2:** Barplot depicting the replication of Figure 2A, using the AmilOrange oligos for overhang base lengths of 0 bp, 1 bp, 2 bp, and 17 bp past the Bsa1 recognition sites.

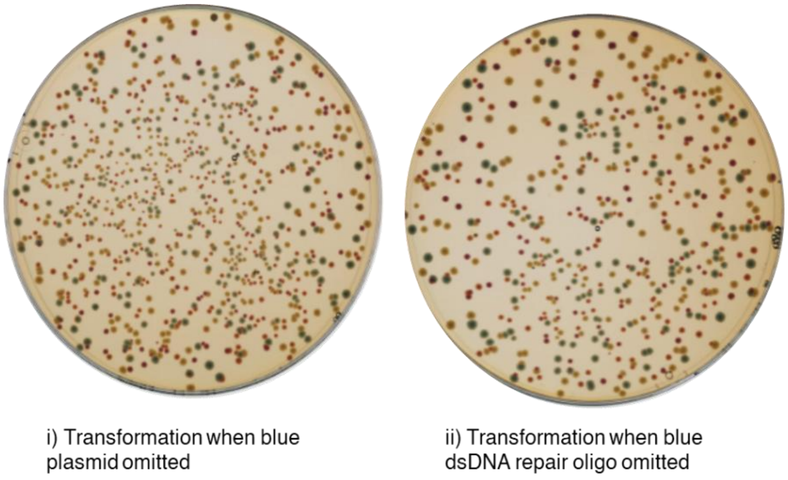


**Figure S3:** Representative agar plates depicting i) the impact of removing the AmilCP (blue) plasmid from the pool of plasmids and ii) the impact of removing the AmilCP (blue) repair oligo from the oPool of oligos. No blue colonies were observed, and there was no increase in white colonies.

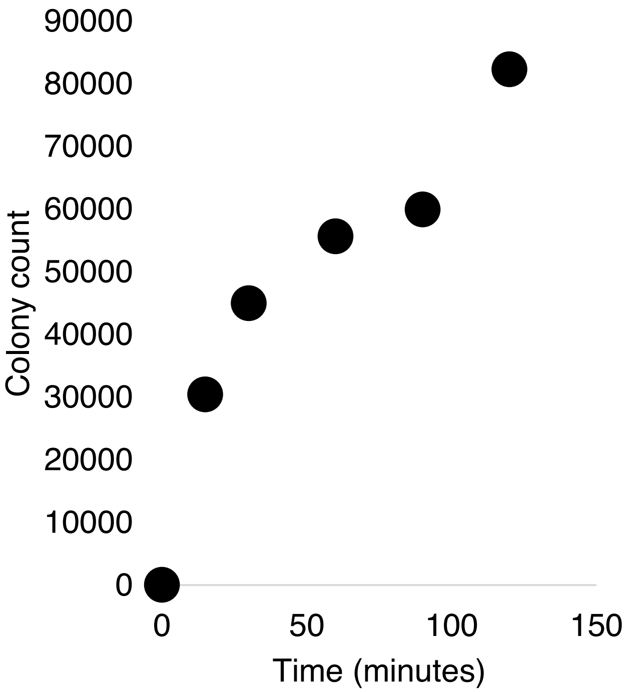


**Figure S4:** Bxb1 reaction kinetics at 37°C observed from transforming the reaction at different times. Reaction after 3 days yielded 126300 colonies (not shown here). Reaction conditions were 50 mM Tris pH 8, 50 mM KCl, 75 mM NaCl, 1 mM EDTA, 100 ug/mL BSA, 5% PEG 8000.

| | | | | | | | | | | | | | | | | |
|---|---|---|---|---|---|---|---|---|---|---|---|---|---|---|---|---|
| [KCl] mM | 100 | 100 | 100 | 100 | 100 | 100 | 80 | 60 | 40 | 20 | 0 | 80 | 60 | 40 | 20 | 0 |
| [NaCl] mM | 100 | 100 | 100 | 100 | 100 | 100 | 80 | 60 | 40 | 20 | 0 | 80 | 60 | 40 | 20 | 0 |
| [Spermidine] mM | 5 | 0 | 2.5 | 5 | 7.5 | 10 | 0 | 0 | 0 | 0 | 0 | 5 | 5 | 5 | 5 | 5 |
| Bxb1 enzyme | - | + | + | + | + | + | + | + | + | + | + | + | + | + | + | + |

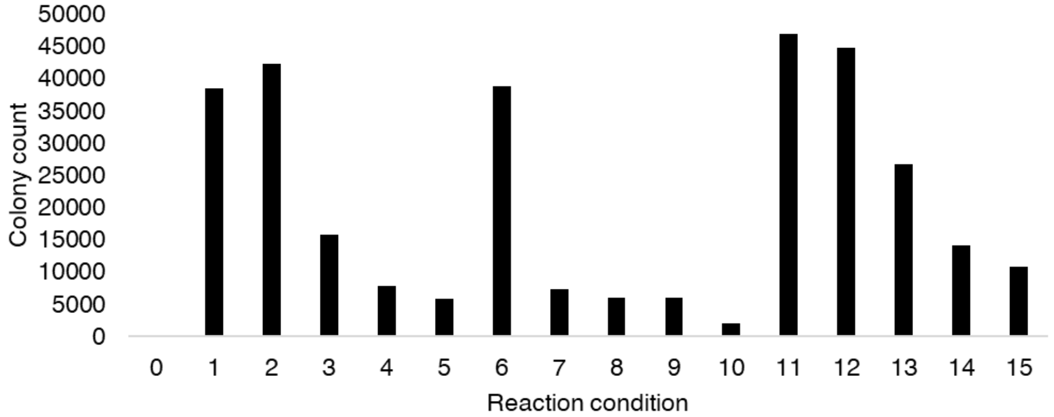


**Figure S5:** Buffer optimization of Bxb1 reaction conditions as assessed by colony count formation after transformation. Reaction time was 60 minutes and temperature was 37°C. Other components of the buffer were 50 mM Tris pH 8, 1 mM EDTA, 100 ug/mL BSA. Reaction table does not include components of the Bxb1 enzyme storage buffer.
